# Mechano-sensitivity of multi-component caveolae

**DOI:** 10.64898/2026.03.03.709391

**Authors:** Niladri Sarkar, Christophe Lamaze, Pierre Sens

## Abstract

Caveolae are small spherical invaginations of the cell membrane with well established mechano-sensing and mechano-regulation functions. The release of caveolae components upon tension elevation is key to their mechao-signalling function. We present a thermodynamic model of caveolae stability under tension based on the phase separation of membrane-associated proteins into invaginated, multi-component membrane domains enriched in the curvature-generating membrane protein caveolin, and stabilised by the curvature-dependent binding of cytosolic proteins: members of the Cavin family which can form a rigid coat over caveolin domains, and the ATPase EHD2 which can form ring-like oligomers at the caveolae neck. This model shows that purely membrane domains show gradual release of their components upon tension elevation. Invaginations stabilised by a Cavin coat can sustain higher tensions with potentially sharp Cavin unbinding at a threshold tension but still release their membrane components smoothly upon tension increase. On the other hand, caveolae stabilised by an EHD2 ring exhibit increased mechano-protection and allow for a sharp release of membrane components. Therefore, the multi-component nature of caveolae self-organisation bestows these membrane domains with a switch-like response to tension variations which leads to the abrupt release of caveolae content beyond a well defined membrane tension threshold.

**Significance statement:** Caveolae are small invaginated domains of Caveolin proteins at the plasma membrane, stabilised by coat of Cavin proteins and a ring of EHD2 proteins at their neck. They contribute to cellular mechano-sensing by disassembling under elevated membrane tension, thereby releasing components that can regulate signalling events. Using a thermodynamic model of protein self-assembly, we show that caveolin domains disassemble smoothly under tension. While the Cavin coat affords mechanical protection without altering this gradual release, the EHD2 ring induces a switch-like release near a threshold tension, enabling acute mechano-sensing.

## Introduction

Caveolae are small (60 − 80 nm) bulb-shaped invagination of the cell membrane that play a major role in the cellular response to mechanical stress by flattening out following an increase of the cell membrane tension (1–3). Caveolae are composed of a membrane domain of the protein caveolin 1 (Cav1), stabilised by the assembly of cytoplasmic Cavins into a coat-like polyhedric structure around the caveola bulb (4– 6), and by the ATP-dependent oligomerisation of the ATPase EHD2 into a ring-like structure at the caveola neck (7, 8), see sketch Fig. 1. Mechano-sensing by caveolae is thought to proceed through the release of caveolae components upon flattening and their involvement in downstream signalling cascades. This include nuclear relocalisation of Cavin (9) and EHD2 (10) released in the cytosol, but also recently discovered direct interaction of freely diffusing Cav1 released at the plasma membrane with signalling effectors (11). To fully appreciate the mechano-signalling role of caveolae, it is essential to determine how their components are released upon changes in membrane tension. This constitutes the focus of the present work.

**Fig. 1.**
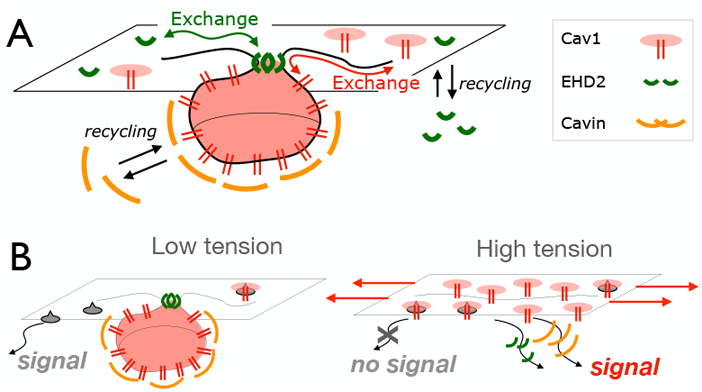
Sketch of the thermodynamic model of multi-component caveolae formation. (A) Cav1 monomers residing on the cell membrane may spontaneously aggregate into invaginated membrane domains in equilibrium with a dilute pool of monomers. The stability of the invaginations is increased by the formation of a Cavin coat binding onto the caveolin domain in a curvature-dependent fashion. Caveolae stability is also enhanced by the formation of an oligomeric ring of EHD2 proteins at their neck, which are in equilibrium with a dilute solution of membrane-bound EHD2 whose concentration is maintained by recycling with a cytoplasmic pool. (B) Possible scenario for caveolae mechano-transduction. An increase of tension destabilizes the caveolar invagination, releasing EHD2 and Cavin in the cytoplasm that may undergo nuclear translocation, and releasing Cav1 monomers in the membrane, that may modify the activity of membrane receptors (here sketched in grey).

Caveolae formation can be understood as a phase separation process driven by intramolecular interactions between components of the caveolar membrane domain and their affinity for a particular membrane curvature (12, 13). Membrane tension increases the cost of membrane deformation, and models based on equilibrium thermodynamics of single component domains have shown that curved domains are stable only below a membrane tension threshold that depends on the concentration of Cav1 at the membrane (12). However, the influence on caveola self-assembly of other cytosolic components binding to the membrane domains—particularly the Cavin coat that covers the caveolar bulb and the EHD2 ring at the caveolar neck— has not been theoretically investigated. Here we propose such a model based on a self-assembly process at thermodynamic equilibrium. We first revisit single-component invaginated membrane domains and show that the disassembly of such structure under tension is gradual, with a progressive decrease of the domain number and size and an increase of the pool freely diffusing Cav1s at the plasma membrane as tension increases. We then study the consequence of assembling a Cavin coat over the curved domain and a ring of EHD2 proteins at the caveola neck. Both provide mechano-protection to caveolae and can exhibit sharp disassembly transition under tension, but the neck ring is essential to permit the abrupt release of membrane components of caveolae above a threshold tension. Our results shed light on the physical role of the different components of caveolae in providing robustness to the function of caveolae as mechano-sensors.

## Model

The shape of membrane domains with spontaneous curvature has been abundantly discussed from a theoretical point of view, but in all cases we are aware of (14–20), this is done varying mechanical parameters for a given domain area rather than letting this area adjust thermodynamically to the mechanical constraints. One notably exception was the case of domains controlled by non-equilibrium fluxes of proteins (21). The present model investigates the thermodynamic equilibrium of the system. It investigates time-scales larger than the typical exchange time between different structures, and it does not discuss the kinetics of their formation, nor potential hysteresis upon dynamic variations of tension(13, 14, 21). Cav1 recycling from the plasma membrane is slow enough so that the total concentration of Cav1 at the membrane remains constant during the self-organisation process (22, 23). On the other hand, the EHD2 and Cavin undergo exchange between membrane-bound and cytosolic states (24, 25) and their chemical potential is assumed to be fixed by the cytosolic pool. The model also relies on the assumption of strong segregation, where the boundary between membrane regions of different composition is sharp and the associated energy cost is characterised by a line tension (14). This is consistent with the strong localisation of Cav1 within the invaginated membrane region (8).

### Thermodynamics of multicomponents caveolae

We consider the situation where a given amount of Cav1 at the cell membrane interacts with a pool of cytoplasmic Cavin and EHD2 proteins to form aggregates of various size and composition. Calling *ρ*_*bud*_ the concentration of domains containing *N*_*c*_ Cav1, *N*_*n*_ EHD2 and *N*_*cc*_ Cavin, and *ρ*_1_ the concentration of freely diffusing Cav1, the free energy per unit area of the system can be written

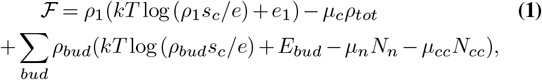

where *kT* is the thermal energy. The first term is the free energy of free Cav1 (monomers), which includes their translational entropy (*s*_*c*_ is the surface area of a monomer) and the energy cost *e*_1_ (*>* 0) of removing a monomer from the caveolin domains. The last term is the free energy of the aggregates, summed over all possible aggregate compositions *bud* = *N*_*c*_, *N*_*n*_, *N*_*cc*_ . It includes their translational entropy, their energy *E*_*bud*_ and the exchange with Cavin and EHD2 reservoirs, of chemical potentials *µ*_*cc*_ and *µ*_*n*_, respectively. These two chemical potentials contain the entropy of the cytosolic proteins and their binding energy to the caveolar membrane. The second term in Eq. 1 introduces the Cav1 chemical potential *µ*_*c*_ as a Lagrange multiplier fixed by the conservation of the total Cav1 concentration *ρ*_*tot*_:

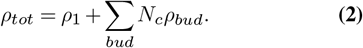

Note that Cav1 monomers usually assemble into small oligomers called 8S even when freely diffusing at the membrane (23, 26), so what we define as Cav1 “monomers” may be thought of as freely diffusing 8S oligomers. Although the mechanism of membrane remodelling by 8S is still debated, their ability to generate membrane curvature is strongly supported by evidences from computer simulation (27–29).

Minimizing the free energy (Eq. 1) with respect to the concentrations of monomers and aggregates yields

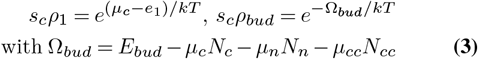

### Energy of a caveola domain

To simplify the model, we restrict the analysis to axisymmetric simple membrane shape of area *S* and uniform curvature *C* (spherical caps). We disregard the membrane tail connecting the cap to the flat membrane, whose contribution to the energy is subdominant (15). We also disregard the possible aggregation of caveolae into superstructure (so-called caveolae rosettes) (30).

Defining the shape parameter *β* = *SC*^2^*/*(16*π*) (*β* = 0 for a flat domain and *β* = 1 for a full sphere), the periphery of the domain is a circle of radius *L* with *πL*^2^ = *S*(1 − *β*). The thermodynamic parameter conjugated to membrane tension is the excess area *Sβ* (the difference between the total and project area of a bud)(14). We assume that Cav1 is densely packed in caveolae, so that *N*_*c*_ = *S/s*_*c*_. Cavin proteins may form a coat on top of the caveolin domain, with *N*_*cc*_ = *ϕ*_*cc*_*S/s*_*cc*_, where *ϕ*_*cc*_ is the Cavin surface fraction in the coat and *s*_*cc*_ is the surface area of one Cavin protein. Similarly the fraction of the invagination’s neck covered with EHD2 proteins is *ϕ*_*n*_ = *N*_*n*_*l*_*n*_*/*(2*πL*), where *l*_*n*_ is the length of one neck protein. One aggregate is thus characterised by four independent parameters, for instance *S, L, ϕ*_*n*_ and *ϕ*_*cc*_. The thermodynamic potential in Eq. 3 may be written as Ω_*bud*_ = Ω_*c*_ +Ω_*n*_ +Ω_*cc*_, with:

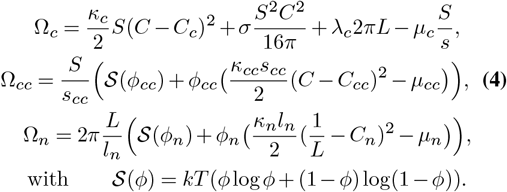

Ω_*c*_ is the contribution of the Cav1 membrane domain, which includes the bending energy (with a bending rigidity *κ*_*c*_ and a preferred curvature *C*_*c*_), the domain line energy (with a line tension *λ*_*c*_) and the work done against the membrane tension *σ*. Ω_*cc*_ is the contribution of the Cavin coat, with a bending rigidity *κ*_*cc*_ and a preferred curvature *C*_*cc*_. Finally, Ω_*n*_ is the contribution of the ring of neck proteins around the caveola neck, assumed to have a preferred (1D) curvature *C*_*n*_ and a (1D) bending rigidity *κ*_*n*_. For the last two contributions, the translational entropy (*ϕ*) of the Cavin and neck proteins within the domain is evaluated using a lattice gas model.

We use the following dimensionless geometric variables

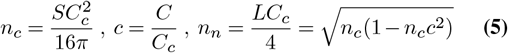

such that *n*_*c*_ = 1 for a full sphere of curvature *C*_*c*_ and 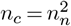 for a flat domain. The shape parameter is *β* = *n*_*c*_*c*^2^.

Minimizing the free energy Ω with respect to the surface fraction *ϕ*_*cc*_ of Cavin on the caveola surface and line fraction *ϕ*_*n*_ of EHD2 at the caveola neck gives:

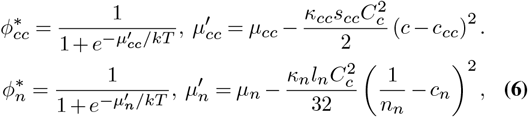

with *c*_*cc*_ = *C*_*cc*_*/C*_*c*_ and *c*_*n*_ = 4*C*_*n*_*/C*_*c*_. This yields the minimized free energy

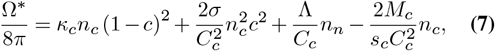

with the effective line tension and chemical potential

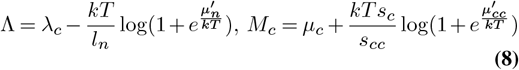

Hence from a thermodynamic point of view, the Cavin coat increases the effective Cav1 chemical potential (thus favouring aggregation) and neck proteins reduces the line tension of the aggregate, both in a shape-dependent fashion.

## Results

In the following, we seek to determine the conditions – in terms of Cav1 concentration and membrane tension – under which Cav1 may self-aggregate into invaginated domains.

The aggregate energy (Eq. 7 with Eq. 8) is minimised for the aggregate shape for a given chemical potential *µ*_*c*_, which is self-consistently computed for a given Cav1 concentration *ρ*_*tot*_ with the conservation relation Eq. 2. The boundaries of the phase diagrams are obtained as the relationship between Cav1 concentration and membrane tension for which the number of Cav1 in aggregates equals the number of freely diffusing Cav1. This is done numerically for a choice of parameter values (see Table S1 in the Supplementary Information - S.I.). Most analytical results given below are obtained for so-called *perfect aggregates* where the bending rigidities 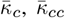 and 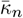 are large and the aggregates, if they form, adopt their preferred values of surface and neck curvatures.

### Single component aggregates

We first consider membrane domains without neck proteins or Cavin coat. We term this “single component” as the physical behaviour is only influenced by the physical properties of the membrane domain (bending rigidity, line tension and spontaneous curvature), although it is clear that such domains are themselves multi-component and contain specific lipid composition (including cholesterol and sphingolipids) in addition to Cav1. The numerical results are presented in Fig. 2. The Cav1 concentration at which aggregates form can be readily identified from the variation of the chemical potential *µ*_*c*_ with the concentration *ρ*_*tot*_ (Fig. 2A). At low concentration, most monomers are freely diffusing and the chemical potential is the one of an ideal gas of monomers *µ*_*c*_ *≃ µ*_*monom*_ = *kT* log *ρ*_*tot*_*s*_*c*_ + *e*_1_. At high concentration, most monomeres are in aggregates with an optimal shape (*S*^*∗*^, *L*^*∗*^) that depends on the membrane tension *σ*. The chemical potential is that of an ideal gas of aggregates: 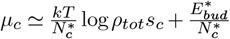, and is almost independent of Cav1 concentration (if 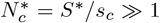). The transition between the two regimes is sharp (Fig. 2A), and occurs at a tension-dependent critical Cav1 concentration 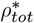 given by 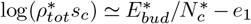. The phase diagram for invaginated domain formation as a function of *ρ*_*tot*_ and *σ* is shown in Fig. 2B. Three possible phases exist: full spheres (*β* = 1), partial spheres shallower than hemispheres (0 *< β <* 1*/*2) and freely diffusing monomers. At low Cav1 concentration, there is a direct transition from full spheres to monomers as the membrane tension increases. At high concentration, there is a first – abrupt – transition from full spheres to hemispheres (*β* ≲ 1*/*2) at a tension value 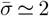, or *σ ≃ λC*_*c*_, followed by a progressive shrinkage of the invaginated domains as the tension increases, and a complete disassembly at a second critical tension (Fig. 2B,C). The amount of freely diffusing monomers increases smoothly with tension during the shrinkage process (Fig. 2D).

**Fig. 2.**
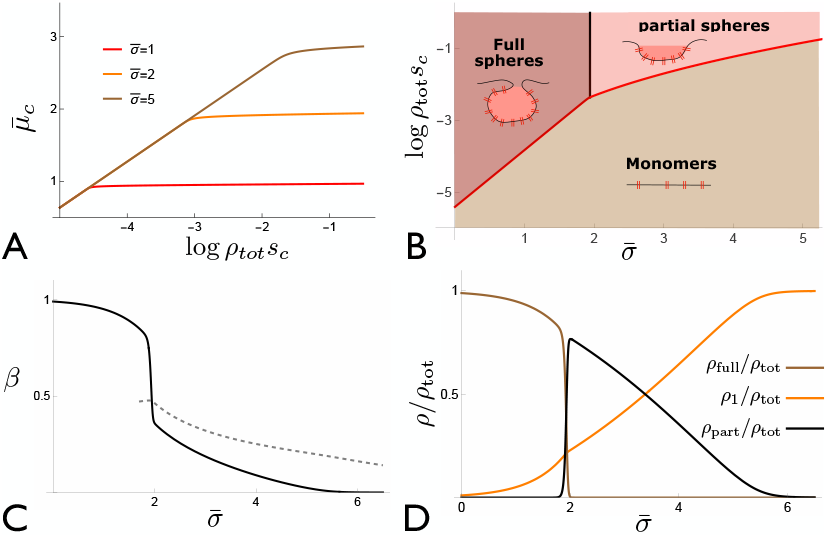
Mechano-sensitivity of single-component aggregates. These results are obtained for perfect (stiff) aggregates with 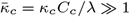 and the dimesionless membrane tension is 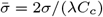. (A) Variation of the Cav1 chemical potential 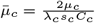 with the log of the Cav1 concentration log *ρ*_*tot*_*s*_*c*_ for different values of the membrane tension, showing a sharp change of slope when domains form. (B) Phase diagram of invaginated domain formation as a function of the total monomer concentration *ρ*_*tot*_ and membrane tension. The three possible phases are monomers (brown), full spherical invaginations (purple) and partial spherical invaginations (pink). (C) Variation of the average shape parameter (sphericity) *β* with tension for a constant *ρ*_*tot*_. The dashed gray line shows the optimal *β* of partial domains. (D) Fraction of the total concentration in full spheres (brown), partial spheres (black), and freely diffusing monomers (orange). For all plots *e*_1_ = 6*kT*, the protein and aggregate energy scales are *e*_0_ *≡ λ*_*c*_*s*_*c*_*C*_*c*_*/*2 = 1.5*kT* and *e*_*N*_ *≡* 8*πλ*_*c*_*/C*_*c*_ = 150*kT* (Table S1). For (C,D), the total concentration is log *ρ*_*tot*_*s*_*c*_ = −1.5.

These results can be understood using the perfect aggregates approximation, corresponding to rigid domains (*κ ≫ λ*_*c*_*/C*_*c*_) which adopt their preferred curvature *C* = *C*_*c*_ regardless of tension. The aggregation transition occurs when the concentration of monomers and aggregates of optimal size are of the same order: *ρ*_1_ *≃ ρ*_*bud*_. From Eq. 3,this occurs when the free energy of the optimal aggregates vanishes (for *N*_*c*_ *≫* 1). The optimal aggregate size *n*_*c*_ and the critical chemical potential *µ*_*c*_ at the transition thus satisfy Ω_*c*_ = 0, 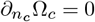. We introduce the dimensionless variables:

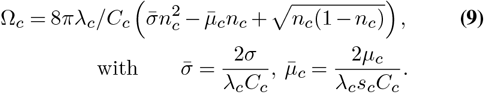

For small tension 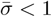, this energy has only two minima, corresponding to monomers (*n*_*c*_ *≃* 0) and full spheres of curvature *C* = *C*_*c*_ (*n*_*c*_ = 1), which coexist when 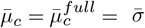 . The energy of any domain coexisting with full spheres (with 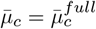) shows a third minimum for hemispheres (*n*_*c*_ = 1*/*2) when 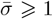, which becomes the global minimum if 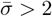. This predicts that if the Cav1 concentration is large enough (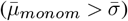), the full spheres are stable if *σ < λ*_*c*_*C*_*c*_. At *σ* = *λ*_*c*_*C*_*c*_, aggregates undergo a transition to hemispheres of the same curvature *C*_*c*_. They loose more area without changing curvature as the tension increases in a way that minimises the free energy Eq. 9. For a given tension, the size and chemical potential of partial spheres satisfy:

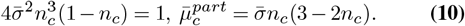

The asymptotic behaviour for large tension (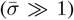) are 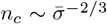 and 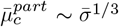 (Eq.S10 in the S.I.). For a given total concentration *ρ*_*tot*_, these aggregates finally dissolve into monomer above a critical tension solution of 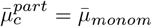, in agreement with the numerical results shown in Fig. 2.

Taking a line tension of order *λ*_*c*_ *≃* 1pN and a spontaneous curvature *C*_*c*_ = 1*/*(25nm) corresponding to *bona fide*, the transition tension tension is within the range of physiological tension: *λ*_*c*_*C*_*c*_ *≃* 4 10^−5^N*/*m. This threshold tension is reduced four-fold if one adopts the spontaneous curvature of Cavin-free Cav1 domains *C*_*c*_ *≃* 1*/*(100nm), indicating a weaker mechanical resistance. This has been proposed before (13), but our model also allows to assess the mechano-sensitivity of the pool of freely diffusing monomers, which was not accessible to the theoretical model of (13) where this pool was assumed constant.

The case of domains with finite bending rigidity (not satisfying 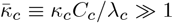) is studied in details in the S.I., and shows no qualitative difference. Full spheres also have the optimal curvature *C*_*c*_. The transition to partial spheres occurs at slightly smaller tension for finite bending rigidity (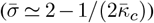) and results in slightly larger and shallower domains. As we expect *κ*_*c*_ *>* 20*kT* (the value for a bare lipid membrane, (31)), we expect 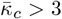, hence a correction of less than 10% on the transition tension for soft domains.

### Caveolae with a Cavin coat: curvature dependent binding

Cavin proteins bind on Cav1-enriched membrane domains and provide structural stability to caveolae (32–34). This is implemented within our equilibrium framework by considering that Cavin binds to Cav1 domains in a curvature-dependent fashion (Ω_*cc*_ in Eq. 4). The Cavin coat has a higher spontaneous curvature than the Cav1 domain (*C*_*cc*_ *> C*_*c*_) (13) and is expected to be more rigid (*κ*_*cc*_ *> κ*_*c*_). Cavin coat formation therefore exhibits cooperativity, since Cavin binding drives the domain curvature closer to the preferred Cavin curvature and promote further binding. This results in an abrupt transition between domains fully coated with Cavin (*ϕ*_*cc*_ *≃* 1) and Cavin-free domains upon membrane tension variation, if the relevant energy scale is larger than the thermal energy *kT* . We show in the S.I. that this occurs if the spontaneous curvature mismatch satisfies 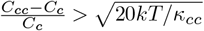 . We concentrate on this situation below and analyse how the presence of a Cavin coat modifies the response of caveolae to variations of membrane tension.

In the limit of stiff domains (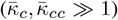), detailed in the S.I. and summarised here, the curvature of Cavin-coated domains with *ϕ*_*cc*_ *≃* 1 is the weighted average of the spontaneous curvature of Cav1 and Cavin: *C*_eff_ = (*κ*_*c*_*C*_*c*_ + *κ*_*cc*_*C*_*cc*_)*/*(*κ*_*c*_ + *κ*_*cc*_). Inserting this into Eq. 8 yields the effective Cav1 chemical potential 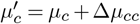, where

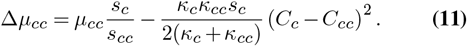

is the net Cavin binding energy, which includes the bending cost due to the spontaneous curvatures mismatch. Cavin-coated domains satisfy the same equilibrium condition as Cavin-free domains (Eq. 10 for partial spheres) with the redefined dimensionless variables 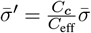 and 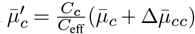, with 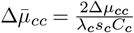 as before.

The phase diagram of caveola equilibrium states as a function of Cav1 and Cavin chemical potentials and membrane tension is shown in Fig. 3A,B. Cavin does not bind to Cav1 domains and the phase diagram is similar to the one-component case Fig. 2B if the net Cavin binding energy 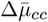 is negative, which means that if the cost of curvature mismatch overcomes the cavin bare binding energy. Full spheres are coated with Cavin if 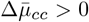, and the transition to partial spheres upon tension increase leads to Cavin unbinding if 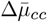 is below a threshold value that increases with *C*_eff_*/C*_*c*_ (it is equal to 1.26 if *C*_eff_ = 2*C*_*c*_). The tension at transition then satisfies 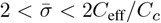. For larger values of 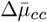 the partial sphere transition occurs at 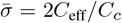 and Cavin remain bound to partial sphere. As the tension increases, cavin progressively unbinds as the Cav1 domain shrink. The mechano-sensitivity of Cavin unbinding is shown in Fig. 3C,and the evolution of the different populations of domains with tension in Fig. 3D. As the net Cavin binding energy 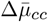 increases, the stability of Cavin-coated partial spheres increases. Consequently, the fraction of freely diffusing Cav1 increases more smoothly with tension (see also Fig.S2).

**Fig. 3.**
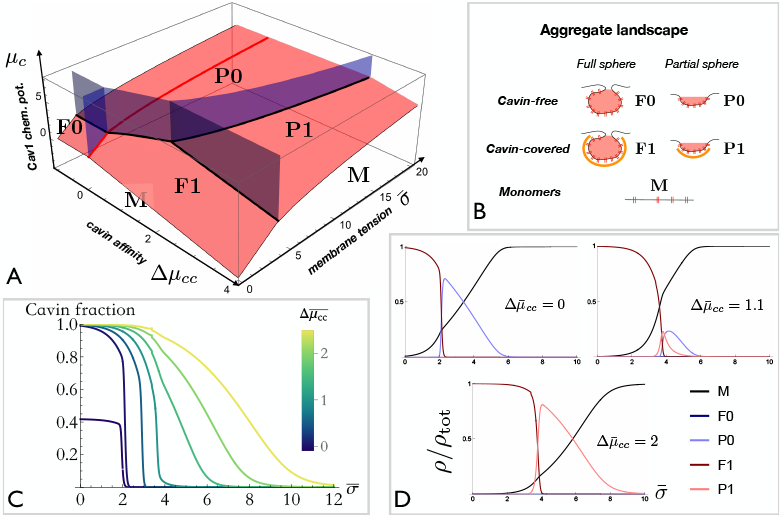
Mechano-sensitivity of caveolae with a Cavin coat. Cav1 and Cavin have different spontaneous curvatures *C*_*c*_ and *C*_*cc*_, and the results are shown in the limit of strongly cooperative Cavin binding (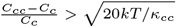, see text). (A) Phase diagram of caveola equilibrium states as a function of Cav1 and Cavin chemical potentials and membrane tension. The phase diagram without Cavin binding (red line, Δ*µ*_*cc*_ *<* 0) corresponds to the phase diagram of Fig. 2B. Five states exist, sketched in (B): full and partial spheres without Cavin (*F* 0 and *P* 0), full and partial spheres with a Cavin coat (*F* 1 and *P* 1), and freely diffusing monomer (*M*). (C) Fraction of Cav1 coated with Cavin as a function of membrane tension. For the lowest Cavin affinity shown (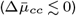), only about half the full spheres are coated with Cavin at low tension. (D) Evolution of the different populations with tension for different 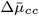. The transition is from Cavin-coated full spheres to Cavin-free partial spheres for intermediate affinity, and to Cavin-coated partial spheres for high Cavin affinity. Parameters are as in Fig. 2 with *C*_eff_*/C*_*c*_ = 2.

To conclude this section, note that tension-mediated transition from Cavin-coated to Cavin-free domains is in principle also possible in the absence of spontaneous curvature mismatch (*C*_*cc*_ = *C*_*c*_) due to a difference of bending rigidity *κ*_*cc*_ ≠ *κ*_*c*_), but is restricted to a very small range of Cavin binding energy, due to the stiffness of Cav1 domains (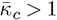, see S.I.)

### Caveolae with an EHD2 neck: neck size-dependent oligomerisation

We now discuss the formation of a ring of EHD2 proteins at the caveola neck. To simplify the discussion, we disregard the possible variation of the Cavin density with tension and assume a Cavin-coated domain with large bending rigidity (“perfect aggregate”). The contribution of neck proteins to the aggregate’s energy is characterised by the effective line tension Λ that depends on the aggregate’s geometry: Eqs.(6,7,8). For perfect aggregates, the energy is: 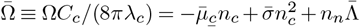, the energy of a spherical cap (Eq. 9) with 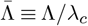 . The EHD2 ring has a preferred curvature *C*_*n*_, or in dimensionless form *c*_*n*_ ≡ 4*C*_*n*_*/C*_*c*_ (Eq. 6).Typical radius and neck radius of caveolae with an EHD2 ring are respectively 2*/C*_*c*_ *≃* 50 nm and 1*/C*_*n*_ *≃* 25 nm (35, 36), which suggests *c*_*n*_ *≃* 4. Note that caveolae with EHD2 have a wider neck than those without (35, 36), which is consistent with our model. Indeed, the neck size in the former case is fixed by the optimal geometry of the EHD2 oligomer: *n*_*n*_ *≃* 1*/c*_*n*_, while it is fixed by line tension in the latter case and is as small as possible. In the limit *n*_*n*_ *≪* 1, the free energy of a perfect aggregate is

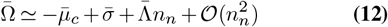

Aggregates with neck proteins are thus predicted to form if 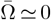, or if 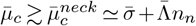. They form preferentially to full spheres if 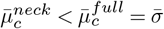, that is if 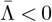.

Under sufficiently large tension, the caveolae neck opens and the ring of neck proteins ring disassembles. Depending on the Cav1 concentration *ρ*_*tot*_, aggregates with neck proteins either break up into smaller invaginations with no neck proteins (characterised by Eq. 10) at a tension solution of 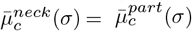, or directly dissolve into monomer at a tension solution of 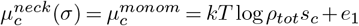, whichever event corresponds to the lowest tension.

The numerical results regarding the mechano-sensitivity of caveolae with a protein ring at their neck are shown Fig. 4 for perfect (stiff) aggregates. This illustrates the two main features provided the ring-like EHD2 oligomer at the caveola neck. The first is the increased mechano-protection against flattening (seen in the phase diagram of Fig. 4A), as observed experimentally (35). The difference between the flattening tension of caveolae with and without neck proteins is 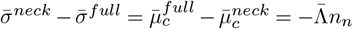. The second is the fact that neck proteins render the tension-triggered release of membrane components of caveolae much more abrupt, as illustrated by the evolution of the different populations of aggregates with tension in Fig. 4B. The diffusing fraction varies as 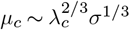 (Eq. 3). For full spheres, *µ*_*c*_ *∼ σ* and the diffusing fraction increases exponentially with the tension. Caveolae without neck proteins first undergo a transition to hemispheres, after which the dependence is much weaker: 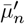 (Eq. 10 with *n*_*c*_ *≪* 1), and the diffusing fraction increases smoothly with tension. The shape transition of caveolae with neck proteins is from a shape close to full sphere (*β≃* 1) to either monomer or very small dimples, depending on *ρ*_*tot*_. The monomer concentration thus exhibits a sigmoidal dependence with tension closer to a switch-like mechano-sensitive response.

**Fig. 4.**
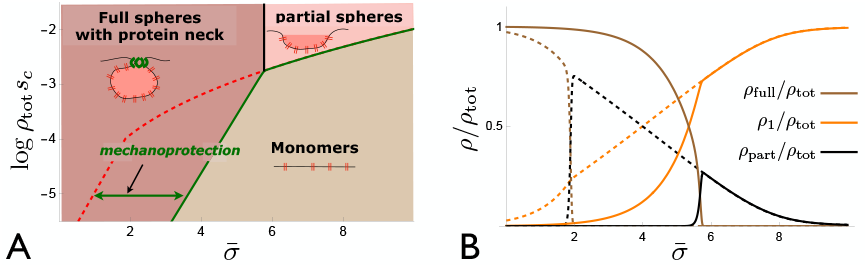
Mechano-sensitivity of caveolae with neck proteins. (A) Phase diagram for the formation of caveolae with a collar of neck proteins (EHD2, sketched in green). The boundary without neck proteins (dashed red line, identical to Fig. 2B) is also shown, highlighting the mechano-protection effect provided by the neck proteins. (B) Fraction of the total Cav1 concentration in full spheres (brown), partial spheres (black), and freely diffusing monomers (orange), with (solid) and without (dashed, as in Fig. 2D) neck proteins. The presence of neck protein renders the release of freely diffusing Cav1 monomers much more abrupt. Parameters are, 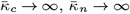 (perfect aggregates), *λ*_*c*_*l*_*n*_*/k*_*B*_*T* = 1.8 and 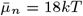 . For B, log *ρ*_*tot*_*s*_*c*_ = −2.4 and *e*_1_ = 6*kT* .

The sharp mechano-sensitive response of caveolae with neck proteins rely on the fact that the EHD2 ring should disassemble as the caveola neck widens during the partial sphere transition. In other words, this requires that the effective neck protein chemical potential 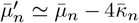 (Eq. 6) becomes negative due to the ring bending energy. For a hemisphere with optimal curvature (*n*_*c*_ = *n*_*n*_ = 1*/*2, with *c*_*n*_ *≃* 4), Eq. 6 gives 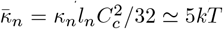, with 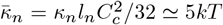 (see Table S1 in the S.I.). Therefore one expects a large shift toward ring disassembly of order 20*kT* per protein for hemispheres. Compare to Cavin unbinding, EDH2 unbinding is therefore insured by the large change of neck curvature at the partial sphere transition.

## Discussion

Caveolae are complexe structures which include several layers of self-organisation. Caveolar membrane domains contain the structural membrane protein Cav1, cholesterol and sphingolipids (37). This peculiar composition endows the domains with high line tension and bending rigidity due to the lipid composition, and a spontaneous curvature due to the asymmetric distribution of Cav1 on the cytoplasmic leaflet of the membrane (13, 31, 38). Theses three physical features could in principle be sufficient to allow for phase separation and the formation of dense and highly curved membrane invaginations (12, 39). Nevertheless, additional supramolecular structures are present in *bona fide* caveolae, including a Cavin coat and a ring of EHD2 proteins at their neck. As several of the caveolae components, including Cavin (9), EHD2 (10) and also freely diffusing Cav1 scaffolds (11) released upon caveolae disruption are involved in various signalling pathways, how these different components are released and made available for mechao-signalling upon membrane tension variations is of fundamental importance. We addressed this issue within the framework of equilibrium thermodynamics.

Our benchmark for mechano-sensation are the so-called single component domains, only involving the membrane protein Cav1 and associated lipid composition. Under vanishing tension, invaginated Cav1 domains can form if the total Cav1 concentration exceeds a threshold given by the domain’s cohesion energy: *µ*_*monom*_ = *kTlogρ*_*tot*_ + *e*_1_ *>* 0. Upon membrane tension increase, domains evolve in one of two ways, depending of a second concentration threshold, equal to the product of the domain line tension by its spontaneous curvature and the protein surface area. For low Cav1 concentration: *µ*_*monom*_ *< λ*_*c*_*s*_*c*_*C*_*c*_, domains remain fully invaginated (“full spheres”) but decrease in number as the tension increases and disappear when *σ > µ*_*monom*_. If *µ*_*monom*_ *> λ*_*c*_*s*_*c*_*C*_*c*_, domains undergo an abrupt transition from fully invaginated to partially invaginated domains (shallower than half-spheres) when *σ ≃ λ*_*c*_*C*_*c*_, then shrink and eventually disappear as the tension further increases. This abrupt shape change is the consequence of the competition between line and surface tension. It bestows membrane domains the ability to regulate membrane tension upon stretch (14), and is reminiscent of a “snap-through” instability of membrane bud under tension that has been described before for domains of constant area (16–18). For appropriate parameter values: *λ*_*c*_ *≃* 1pN and *C*_*c*_ *≃* 1*/*(100nm) (for Cavin-free domains), the critical membrane tension (*λ*_*c*_*C*_*c*_ *≃* 10^−5^N*/*m) is in the low range of physiological tensions (40). This could explains why only partially invaginated Cav1 domains –so-called dolines– have been observed in the absence of Cavin (13, 41, 42). Remarkably, their presence at finite tension indicates that the concentration of Cav1 is sufficient for fully invaginated Cavin-free domains to form if the cell tension is lowered, for instance by subjecting the cell to an hyper-osmotic shock.

Importantly, the amount of freely diffusing Cav1 released by single-component domains increases gradually with tension, as the domains decrease in number and shrink (Fig. 2). Thus, while a mechano-signaling pathway based on their availability is certainly conceivable, it lacks a switch-like character. Can Cavin proteins, which assemble into a coat complex on caveolin domains after recruiting from the cytosol, influence the mechanical response of these domains? The cavin coat is expected to increase the domain’s rigidity, but order of magnitude estimates show that this has little effect. Cavin is also expected to increase the domain’s curvature, which directly impact its mechano-resistance through the threshold tension *λ*_*c*_*C*_*c*_. Cavin-coated domain should be fully invaginated up to tension level (*≃* 4 *×* 10^−5^N*/*m) above the resting tension in many mammalian cells. however, the Cavin coat does not affect the gradual release of membrane component upon tension elevation, which remains similar to the one component case (Fig. 3). Conversely, the release of cytosolic components (the Cavins) can occur abruptly, since Cavin binding and unbinding is a cooperative process: the binding of one Cavin modifies the domain curvature in a way that promotes further Cavin binding. When the Cavin binding energy to the Cav1 domain is high, Cavins remain associated with partially invaginated domains and are released gradually, alongside other membrane components. In contrast when the binding energy is moderate, Cavins detach abruptly at the threshold tension where the domain transitions from fully invaginated to partially invaginated (Fig. 3). This appears to be a good strategy to create a mechano-sensitive switch based on abrupt release of Cavin into the cytosol at a relatively sharp tension threshold (9, 43). Whether this corresponds to the situation *in vivo* is not yet clear. Partially invaginated or even flat domains coated with Cavin have been observed by electron microscopy (42, 44), but might correspond to transient structures fixated en route to formation or disassembly.

A fully mature caveola generally possesses a ring of EHD2 proteins at its neck. This ring is equivalent to a curvature-dependent line tension (Eq. 8), which favours a neck of particular curvature (slightly larger than the one without EHD2 (35)) with a particular rigidity. The ring provides mechanoprotection to the invaginated structure, and disassembles at a higher tension, corresponding to the mechanical stress required to overcomes the EHD2 binding strength, in agreement with experimental observations (35). With the parameters used in Fig. 4 the critical tension is three times higher with EHD2 than without. Ring disassembly and EHD2 release into the cytosol is predicted to be abrupt, which concurs with experimental observations that EHD2 is not found in smaller Cav1 structures (45), so that EHD2 could also be involved in a switch-like mechano-sensitive pathway. Importantly, the release of freely diffusing Cav1 scaffolds also occurs in a sigmoid fashion (Fig. 4), and could trigger a sharp mechano-sensitive cell response based on the interaction of free Cav1 with signalling effectors (11).

Experimental observations on the size distribution of caveolae are scarce. Recent observations suggest an equal amount of small Cav1 scaffolds (considered as monomers in our model) and larger, partially invaginated domains (partial spheres in our model) in the absence of Cavin, while about half of the domains turn into fully formed caveolae (full spheres in our model) in the presence of Cavin (41, 42). This suggests that the conditions of tension and Cav1 concentration in this particular experimental setup are close to the partial sphere-monomer transition for domains without Cavin, and close to the full sphere-partial sphere-monomer transition with Cavin. This is consistent with our predictions provided that: (i) the full sphere to partial sphere transition tension (*λ*_*c*_*C*_*c*_) is low for domains without Cavin, possibly because of a small spontaneous curvature, which suggests a relatively weak mechano-sensitivity of these structures, (ii) domains with Cavin are in a region of the phase diagram close to the “triple point” where full spheres, partial spheres and monomers coexist. Interestingly, this also happens to be the region of the phase diagram where mechano-sensitivity is the strongest.

In summary, Cav1 membrane domains are predicted to release their components gradually upon tension increase. The Cavin coat increases the mechanical stability of the invaginated domain, and its release in the cytosol upon tension increase can be expected to be abrupt and could trigger a sharp mechano-sensitive response, but the relase of memrbane components remains gradual. The EHD2 ring also stabilises the structure and gets released abruptly upon tension icnrease, but also permits a much sharper release of membrane components (including Cav1) which can then trigger a sharp mechano-sensitive response at the cell membrane. Importantly, this sharp release extends to any component of Cav1-enriched domains, including ion channels are preferentially located in caveolae (46, 47) but also components not directly coupled to membrane mechanics.

## ACKNOWLEDGEMENTS

We acknowledge stimulating discussion with Cédric Blouin (Institut Curie) and Stéphane Vassilopoulos (Institut de Myologie, Paris). This work was supported by the French National Research Agency (DECAV-RECAV, ANR-14-CE09-0008-03 (NS, CL PS), MOTICAV ANR-17-CE13-0020-01 (CL PS) and CAV-SM ANR-20-CE13-0002-01(CL)), the Labex Cell(n)Scale (ANR-10-LBX-0038) part of the IDEX PSL (ANR-10-IDEX-0001-02 PSL) (CL, PS), the Cell and Tissue Imaging core facility (PICT-IBiSA), member of the France-BioImaging national research infrastructure, supported by Fondation ARC pour la recherche sur le cancer (Programme Labellisé PGA1-RF20170205456), CEFIPRA, project 6301-1 (CL) and the European Union, ERC, PushingCell, project 101071793 (PS)

